# Structure and methyl-lysine binding selectivity of the HUSH complex subunit MPP8

**DOI:** 10.1101/2023.12.21.572340

**Authors:** Nikos Nikolopoulos, Shun-ichiro Oda, Daniil M. Prigozhin, Yorgo Modis

## Abstract

The Human Silencing Hub (HUSH) guards the genome from the pathogenic effects of retroelement expression. Composed of MPP8, TASOR, and Periphilin-1, HUSH recognizes actively transcribed retrotransposed sequences by the presence of long (>1.5-kb) nascent transcripts without introns. HUSH recruits effectors that alter chromatin structure, degrade transcripts, and deposit transcriptionally repressive epigenetic marks. Here, we report the crystal structure of the C-terminal domain (CTD) of MPP8 necessary for HUSH activity. The MPP8 CTD consists of five ankyrin repeats followed by a domain with structural homology to the PINIT domains of Siz/PIAS-family SUMO E3 ligases. AlphaFold-Multimer modeling predicts that the MPP8 CTD forms extended interaction interfaces with a SPOC domain and a domain with a novel fold in TASOR. The MPP8 chromodomain, known to bind the repressive mark H3K9me3, binds with similar or higher affinity to sequences in the H3K9 methyltransferase subunits SETDB1, ATF7IP, G9a, and GLP. Hence, MPP8 promotes heterochromatinization by recruiting H3K9 methyltransferases. Our work identifies novel structural elements in MPP8 required for HUSH complex assembly and silencing, thereby fulfilling vital functions in controlling retrotransposons.

## Introduction

Transposable elements (TEs) have been coevolving with the genomes of all organisms since the early stages of life. Some TEs, such as long interspersed nuclear elements (LINEs), are thought to have evolved in early eukaryotes (Goodier, 2016). TEs can also be acquired when viral DNA integrates into the genome of a host germline cell. Why TEs are so ubiquitous and abundant, accounting for more than half of the human genome (Friedli & Trono, 2015), is not fully understood. TEs are known to serve as a genetic reservoir from which new genes, regulatory elements, and transcriptional networks can emerge (Chuong *et al*, 2016; Dupressoir *et al*, 2012; Friedli & Trono, 2015; Pastuzyn *et al*, 2018; Zhou *et al*, 2004). However, inappropriate expression of TEs has been associated with various genetic diseases including autoimmune diseases, neurodegeneration, hemophilia, cystic fibrosis, and cancer (Goodier, 2016; Hancks & Kazazian, 2016; Hung *et al*, 2015; Lamprecht *et al*, 2010). The primary mechanism cells have evolved to control TE expression is epigenetic transcriptional silencing. Transcriptional repression of TEs is particularly important in embryogenesis, chronic infection, and stress responses, when epigenetic modifications are more pro-transcriptional (Azebi *et al*, 2019). In vertebrates, the Human Silencing Hub (HUSH) is a key repressor of retrotransposons, the subset of TEs that can replicate by retrotransposition (Tchasovnikarova *et al*, 2015). The hundreds of genomic loci repressed by the HUSH complex include recently integrated viral elements, LINE-1s, and a subset of endogenous genes generated through retrotransposition of cellular mRNAs (Douse *et al*, 2020; Liu *et al*, 2018; Seczynska *et al*, 2022; Tchasovnikarova *et al*., 2015). HUSH target loci have in common that they encode transcripts longer than 1.5 kb, are transcriptionally active, and are products of retrotransposition and therefore lack introns (Seczynska *et al*., 2022).

The HUSH complex consists of three proteins: TASOR (Transgene activation suppressor), MPP8 (M-phase phosphoprotein 8), and Periphilin (PPLHN1, isoform 2) (Tchasovnikarova *et al*., 2015). Periphilin targets HUSH to its genomic loci by binding to nascent transcripts with exons longer than 1.5 kb (Seczynska *et al*., 2022). Periphilin binds nascent RNA with a disordered, arginine/tyrosine-rich N-terminal domain with self-aggregating properties (Prigozhin *et al*, 2020). MPP8 was recently reported to contribute to HUSH targeting by binding to termination factor WDR82 that is enriched at sites with high RNA polymerase II occupancy, which include long exons (Spencley *et al*, 2023). TASOR binds both Periphilin and MPP8 to form the HUSH complex (Prigozhin *et al*., 2020; Tchasovnikarova *et al*., 2015). TASOR contains three folded domains required for HUSH activity: a pseudo-PARP, a predicted SPOC (Spen Paralogue and Orthologue C-terminal) domain, and a domain of unknown fold, DomI (Douse *et al*., 2020; Tchasovnikarova *et al*., 2015). The pseudo-PARP binds weakly to ssRNA and is required for HUSH activity, but how this domain contributes to gene silencing remains unclear (Douse *et al*., 2020). A structurally homologous pseudo-PARP in TEX15, a piRNA-dependent transposon silencing factor in the male germline, can functionally substitute for the pseudo-PARP of TASOR (Schopp *et al*, 2023). MPP8 recruits ATF7IP (activating transcription factor 7 interacting protein), an obligate binding partner and stabilizing factor of the histone H3 lysine 9 (H3K9) methyltransferase SETDB1 (SET domain bifurcated histone lysine methyltransferase 1) (Timms *et al*, 2016). HUSH silencing additionally requires MORC2 (microrchidia family CW-type zinc finger 2), a DNA-binding ATPase thought to be a chromatin remodeler (Douse *et al*, 2018; Tchasovnikarova *et al*, 2017). Thus, in the current model of HUSH-dependent repression, Periphilin binds to nascent transcripts of newly retrotransposed sequences, MPP8 recruits SETDB1/ATF7IP to deposit the transcriptionally repressive H3K9me3 mark, and MORC2 is recruited to promote chromatin compaction (**Figure 1A**). Notably, a subset of MPP8 chromatin binding sites lack TASOR and Periphilin, suggesting a HUSH-independent function of MPP8 chromatin binding (Spencley *et al*., 2023). Consistent with this, MPP8 recruits the NEXT (nuclear exosome targeting) complex to chromatin (via an interaction with ZCCHC8), leading to exosome-dependent decay of non-polyadenylated TE transcripts, whereas HUSH represses polyadenylated transcripts (Garland *et al*, 2022). Together, the transcriptional repression activities of MPP8 protect the DNA-hypomethylated pluripotent ground state of stem cells (Muller *et al*, 2021). MPP8 depletion causes transcription of RNAs that induce interferon and DNA damage responses, due to recognition by the RNA sensor MDA5 and increased retrotransposition activity, respectively (Gu *et al*, 2021; Tunbak *et al*, 2020). These responses are thought to account for inhibition of acute myeloid leukemia and sensitization of tumor cells to checkpoint blockade upon MPP8 knockout (Griffin *et al*, 2021; Gu *et al*., 2021).

**Fig. 1.**
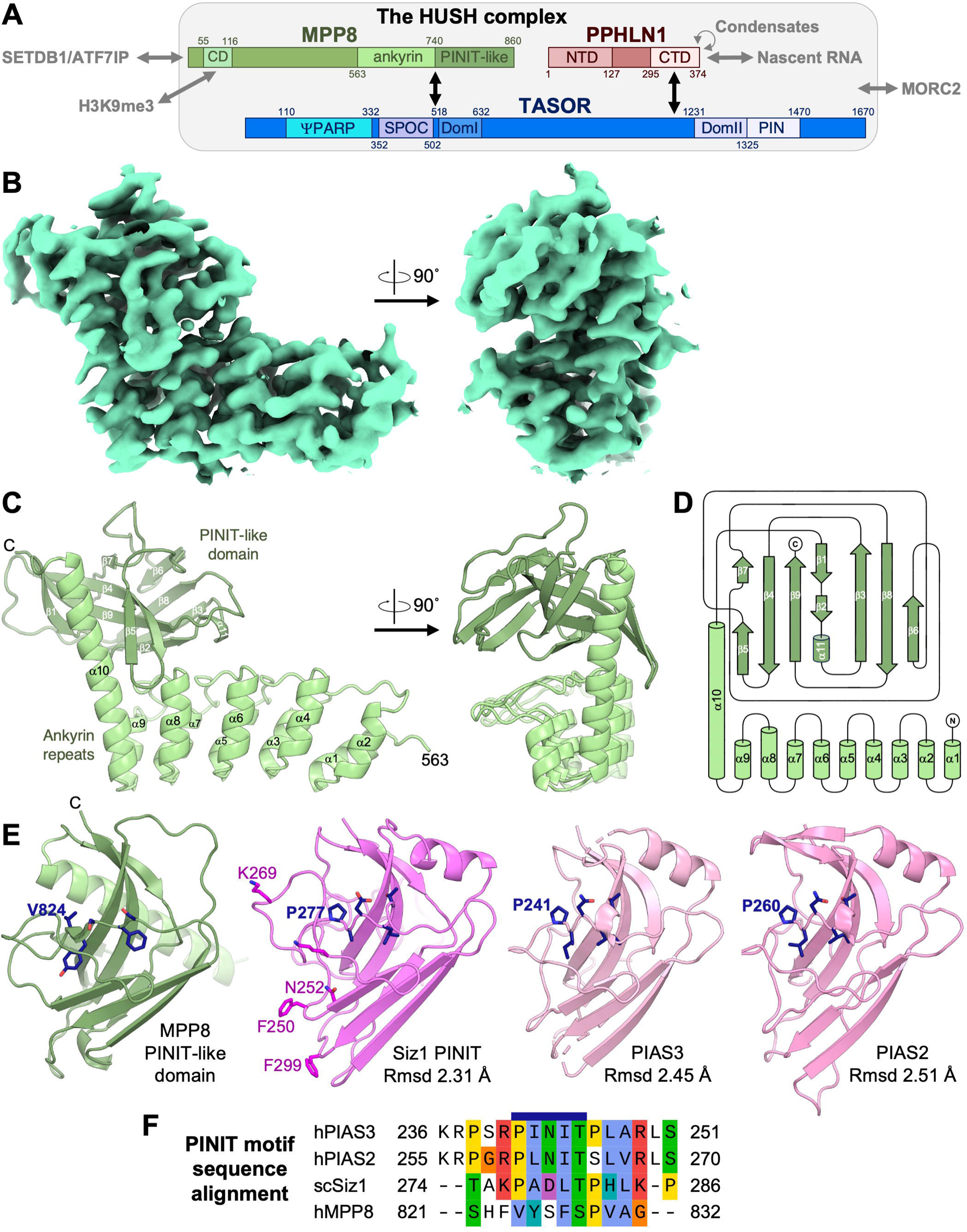
Crystal structure of the MPP8 C-terminal domain (CTD) and structural homology to PINIT domains. (**A**) Domain diagram of the HUSH complex (drawn to scale) and its interactions. (**B**) Weighted 2*F*_o_ – *F*_c_ electron density map for one MPP8 CTD subunit. Created with ChimeraX (Pettersen *et al*, 2021) using contour level 0.104. See **Table 1** for crystallographic data collection, refinement, and validation statistics. (**C**) Crystal structure of the MPP8 CTD. See **Fig. S1A** for representative electron density snapshots and non-crystallographic symmetry with disulfide-bonded dimers. (**D**) Secondary structure topology of the MPP8 CTD. Created with TopDraw (Bond, 2003). (**E**) Structural homology of the MPP8 β-sandwich to the PINIT domains from Siz/PIAS-family E3 SUMO ligases *Saccharomyces cerevisiae* Siz1 (Yunus & Lima, 2009), human PIAS2 (PDB 4FO9), and human PIAS3 (PDB 4MVT). Rmsd, root mean square deviation between atoms in a superposition with MPP8. (**F**) Multiple sequence alignment of the PINIT motif (blue bar) and flanking sequences of PIAS3, PIAS2, Siz1 with the MPP8 sequence aligned based on structural superpositions.

**Table 1.** Crystallographic data collection, refinement, and validation statistics for MPP8 CTD.

|  |  |
| --- | --- |
| Space group | *P* 2<sub>1</sub> 2<sub>1</sub> 2<sub>1</sub> |
| Cell dimensions |  |
| *a*, *b*, *c* (Å) | 102.56 179.53 228.08 |
| $\alpha$, $\beta$, $\gamma$ (°) | 90, 90, 90 |
| Resolution (Å) | 93.5 – 3.0 (3.107 – 3.0) |
| Total reflections | 1,097,353 (94,635) |
| Unique reflections | 84,956 (8,368) |
| Multiplicity | 12.9 (11.3) |
| Completeness | 98.6 (94.2) |
| / $\sigma(I)$ | 14.36 (1.46) |
| Wilson *B*-factor | 99.7 |
| *R*<sub>merge</sub> | 0.167 (3.20) |
| *R*<sub>pim</sub> | 0.0482 (0.979) |
| CC1/2 | 0.999 (0.266) |
| **Refinement and Validation** |  |
| Resolution (Å) | 93.5 – 3.04 (3.15 – 3.04) |
| No. reflections, working set | 80,744 (7,731) |
| No. reflections, test set | 4,133 (423) |
| *R*<sub>work</sub> | 0.2554 (0.5028) |
| *R*<sub>free</sub> | 0.2948 (0.5530) |
| No. non-hydrogen atoms | 17,740 |
| No. ligand or solvent atoms | 0 |
| *No. MPP8 chains per* | *8* |
| *a.s.u.* |  |
| *Average B-factor* | *131* |
| *RMSD* |  |
| *Bond lengths (Å)* | *0.005* |
| *Bond angles (°)* | *0.85* |
| *Ramachandran plot* |  |
| *Favored (%)* | *97.8* |
| *Allowed (%)* | *2.2* |
| *Outliers (%)* | *0.0* |
| *Clashscore (Phenix 1.20.1)* | *12.3* |
| *PDB accession code* | *8QFB* |

*Numbers in parentheses are for the highest resolution shell*

Biochemical and structural data for the protein domains essential for HUSH function are sparse. This study focuses on delineating the structural and biochemical properties of MPP8 and how they contribute to transcriptional repression. MPP8 contains an N-terminal chromodomain (CD, residues 59-118) and a set of five predicted ankyrin repeats (Douse *et al*., 2020). The predicted ankyrin repeats (residues 548-728) and the following C-terminal region (residues 729-860) are both absolutely required for binding to TASOR and for HUSH repression of transgenes and LINE-1s (Douse *et al*., 2020; Muller *et al*., 2021). In contrast, HUSH repression of an integrated lentivirus reporter is maintained, although slightly delayed, upon deletion of the CD, or indeed the entire N-terminal region of MPP8 (residues 1-499) (Douse *et al*., 2020). The MPP8 CD binds di- or trimethylated H3K9 peptides with micromolar dissociation constants (Bua *et al*, 2009; Chang *et al*, 2011a; Quinn *et al*, 2010). A crystal structure of the MPP8 CD bound to the H3K9me3 peptide showed that the trimethyl-lysine is bound in a cage formed by one acidic and three aromatic residues (Chang *et al*., 2011a). Binding to mono- or unmethylated H3K9 peptides, and to certain methylated or unmethylated H3K4 and H4K20 peptides, is 20- to 600-fold weaker but still measurable (Chang *et al*., 2011a; Quinn *et al*., 2010).

The relatively low affinities and methylation specificities of MPP8 CD for lysine-methylated peptides, and its auxiliary role in HUSH-dependent lentivirus repression, raise the question of whether this domain has other binding partners or functions. We note that H3K9 mimic sequences have been identified in all four protein components of two of the most important mammalian H3K9 methyltransferase complexes, G9a/GLP and SETDB1/ATF7IP (Chin *et al*, 2007; Sampath *et al*, 2007; Tsusaka *et al*, 2018). G9a/GLP automethylates the H3K9 mimic sequence in G9a, which can then bind the CD of HP1 (Chin *et al*., 2007; Sampath *et al*., 2007). G9a/GLP also methylates the H3K9 mimic sequence of ATF7IP at Lys16 (Tsusaka *et al*., 2018). ATF7IP methylated at Lys16 was found to co-immunoprecipitate with the CD of MPP8 (Tsusaka *et al*., 2018). Mutation or deletion of ATF7IP Lys16 abrogated MPP8 chromatin binding and delayed HUSH-dependent repression of a reporter transgene in mouse embryonic stem cells (Tsusaka *et al*., 2018), a similar phenotype to deletion of the MPP8 CD or mutation of its H3K9me3 binding motif (Douse *et al*., 2020). Similarly, MPP8 was found to co-immunoprecipitate with GLP and bind Dnmt3a via H3K9-like sequences, which are both methylated by G9a/GLP (Chang *et al*, 2011b; Kokura *et al*, 2010). Thus, MPP8 may bind to H3K9-like peptides in a broader set of proteins. However, only the interactions of the MPP8 CD with H3K9me3 and Dnmt3aK47me2 have been validated biochemically using purified components (Chang *et al*., 2011a; Chang *et al*., 2011b).

Here, we report the crystal structure of the C-terminal domain (CTD) of MPP8 necessary and sufficient for HUSH activity consisting of an ankyrin repeat domain and a β-sandwich with a PINIT domain fold. We used AlphaFold-Multimer to generate a high-confidence model of the MPP8-TASOR complex, which suggests that the MPP8 CTD binds to a SPOC domain and a domain with a novel fold, DomI, in TASOR. The AlphaFold model of the MPP8-TASOR complex is experimentally supported by HUSH activity assays with truncations of MPP8 and TASOR (Douse *et al*., 2020). Isothermal titration calorimetry measurements show that the MPP8 CD binds to H3K9-like peptides from G9a/GLP and SETDB1/ATF7IP with similar or higher affinities to H3K9me3. In the context of the previously reported phenotypes of MPP8 deletions, we propose that these interactions accelerate or amplify transcriptional repression through increased recruitment of H3K9 methyltransferases in a read/write mechanism. Our work identifies novel structural elements in MPP8 required for HUSH assembly and effector recruitment.

## Results

### Structure of MPP8 CTD reveals ankyrin repeats and a PINIT-like domain

The region of MPP8 required for transcriptional repression of an integrating lentiviral reporter has been mapped to within residues 500-860 (Douse *et al*., 2020). Moreover, residues 528-860 of MPP8 are sufficient for HUSH complex formation (Douse *et al*., 2020). Having established the functional importance of the MPP8 C-terminal domain (CTD), we sought to determine its structure to gain insights on how it contributes to gene silencing as part of the HUSH complex. A 3.04 Å resolution crystal structure of MPP8 CTD (residues 563-860) was determined by molecular replacement using the AlphaFold prediction for the CTD as the search model (**Fig. 1B**, **Table 1** and **Fig. S1A**). The crystal structure reveals a bipartite fold consisting of five ankyrin (helix-loop-helix) repeats followed by a β-sandwich domain (**Fig. 1C,D**). The last α-helix of the ankyrin repeats, helix α10, is twice as long as the other ankyrin-repeat helices and extends along the β-sandwich domain (**Fig. 1C,D**). The β-sandwich is tightly packed onto the ankyrin repeats and helix α10. One of the β-sheets (strands β1, β4, β5 and β9) wraps around the C-terminal half of helix α10. The β-sandwich forms additional contacts with the loops of the last two ankyrin repeats (**Fig. 1C,D**).

A structural similarity search with DALI (Holm & Rosenstrom, 2010) revealed structural similarity of the MPP8 β-sandwich domain (residues 741-860) to the PINIT domains of the Siz/PIAS RING family of SUMO E3 ligases (Duval *et al*, 2003). The most similar structures were those of the PINIT domains from *Saccharomyces cerevisiae* Siz1 (Yunus & Lima, 2009), human PIAS3, and human PIAS2 (**Fig. 1E**). The domains have the same topology and overall fold (2.3-2.5 Å RMSD, Z-score 8.4-9.4), although the Siz/PIAS PINIT domains have a flexible N-terminal linker in place of the rigid helical linker (helix α10) in MPP8. The PINIT domain is the substrate recognition domain of Siz/PIAS SUMO E3 ligases and is required for conjugation of SUMO to their substrate proteins (Streich & Lima, 2016; Yunus & Lima, 2009). The PINIT domain of Siz1 is required for SUMOylation of splicing factor Prp45 and the proliferating cell nuclear antigen (PCNA) in yeast (Pfander *et al*, 2005; Streich & Lima, 2016). In humans, PIAS2 and PIAS3 have been reported to SUMOylate various substrates including Aurora-B kinase, Argonaute-2, and the INO80 chromatin remodeling complex (Ban *et al*, 2011; Cox *et al*, 2017; Josa-Prado *et al*, 2015; Salas-Lloret *et al*, 2023). The PINIT amino acid sequence motif is not conserved in the PINIT-like domain of MPP8 (**Fig. 1F**). Indeed, there is no detectable sequence similarity between the PINIT domains of MPP8 and Siz/PIAS. MPP8 also lacks a RING domain and is therefore not an E3 ligase. The presence of a PINIT-like domain in MPP8 remains notable given that HUSH subunits TASOR and Periphilin, and HUSH effectors SETDB1, ATF7IP and MORC2 all contain SUMO acceptor sites (Hendriks *et al*, 2017).

During refinement of the crystal structure of MPP8 CTD, we noticed that the eight subunits in the asymmetric unit form four disulfide-linked homodimers. The intermolecular disulfide bond linking the homodimers was formed by cysteine 799, located in the loop between strands β5 and β6 of the PINIT-like domain (**Fig. S1B**). Moreover, 10-15% of the recombinant MPP8 CTD protein migrated as a dimer in size-exclusion chromatography (SEC), indicating that some dimers formed in solution despite the presence of reducing agent (1 mM DTT or 0.5 mM TCEP) in the buffer (**Fig. S2A**). However, analysis with the PISA server (39) showed that the dimers have a small interface area (377 Å) and no strong contacts other than the disulfide bond. We therefore conclude that in the context of the reducing intracellular environment the disulfide bond and dimer interface observed in the crystal are unlikely to be physiologically relevant.

### Molecular modeling of the MPP8-TASOR complex with AlphaFold-Multimer

We showed previously using purified recombinant proteins that MPP8 CTD forms a 1:1 complex with a TASOR fragment (residues 354-633) that was predicted to contain two folded domains: a SPOC domain and DomI, a domain of unknown structure and function (Douse *et al*., 2020). We showed that DomI was required for co-immunoprecipitation with MPP8, and deletion of either the SPOC or DomI domains resulted in loss of HUSH-dependent transgene reporter repression (Douse *et al*., 2020). As we were unable to crystallize or obtain a cryo-EM reconstruction of an MPP8-TASOR complex, we generated a predicted atomic model with AlphaFold-Multimer (Evans *et al*, 2021; Jumper *et al*, 2021; Mirdita *et al*, 2022), as implemented by ColabFold (Mirdita *et al*., 2022). With only the sequences of human TASOR and MPP8 as the input, AlphaFold generated 1:1 MPP8-TASOR complexes with high confidence scores (average pLDDT = 88-90) and low predicted aligned error (**Fig. 2A**). All the generated AlphaFold models had the same overall structure, with or without the use of model templates from the Protein Data Bank (PDB). Residues 125-535 of MPP8 were consistently unstructured in AlphaFold predictions, as were amino acids flanking the SPOC and DomI domains of TASOR. Moreover, the pseudo-PARP domain of TASOR did not interact with MPP8 in the predicted models. The boundaries of structural disorder in the AlphaFold-Multimer predictions suggest that the core complex consists of MPP8 CTD residues 536-860 and TASOR SPOC-DomI residues 352-632 (**Fig. 2A** and **Fig. S3C**).

**Fig. 2.**
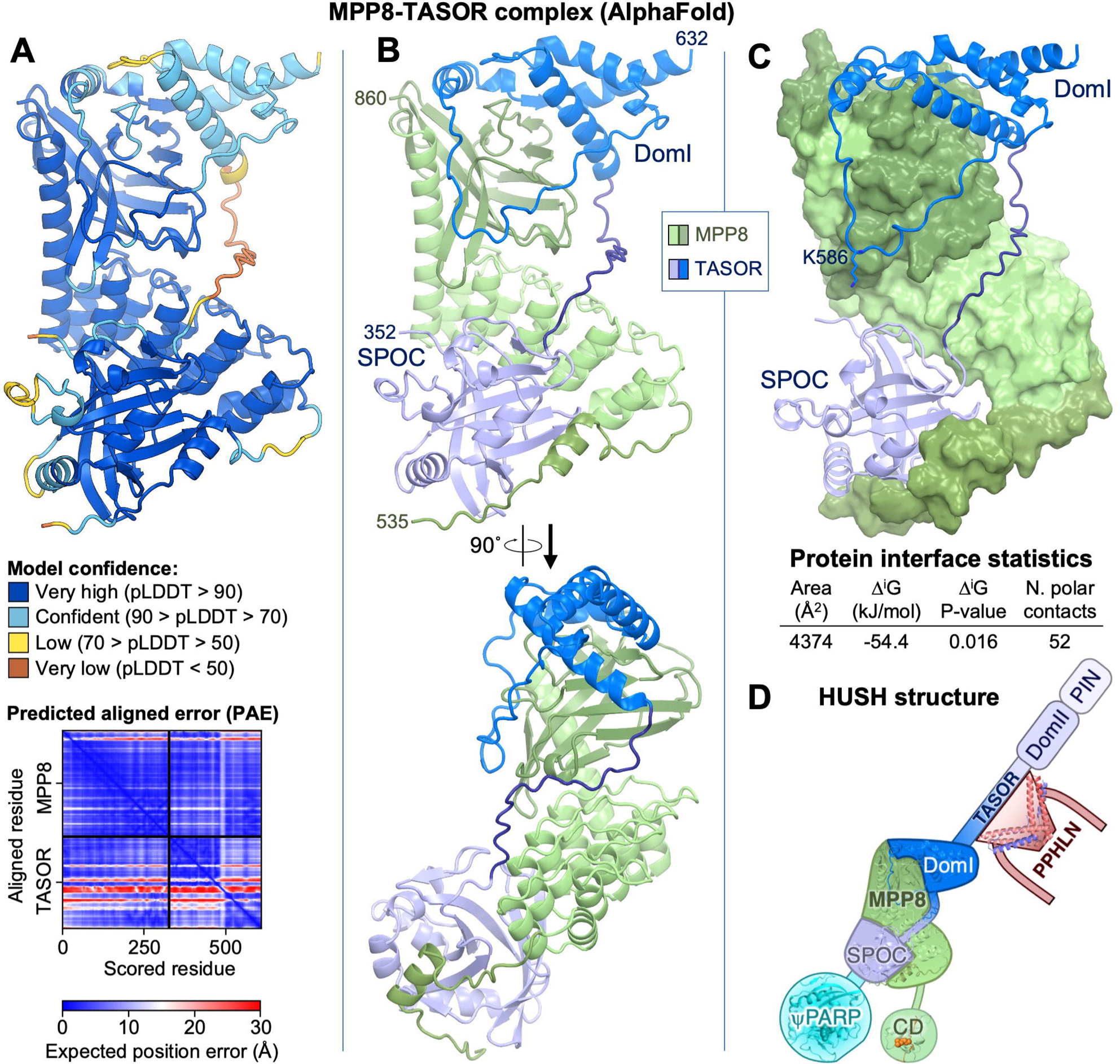
Structural modeling of the MPP8-TASOR complex with AlphaFold-Multimer. (**A**) AlphaFold model of the MPP8-TASOR complex colored by model confidence (pLDDT, according to the color code shown). Lower panel: predicted aligned error plot. (**B**) AlphaFold model of the MPP8-TASOR complex colored by domain. (**C**) AlphaFold model of the MPP8-TASOR complex with MPP8 in surface representation and with PISA protein interface parameters. See **Fig. S3** for closeups of the MPP8-TASOR interface; a comparison of the intermolecular contacts formed by the PINIT domains of MPP8 and Siz1; and a comparison of the TASOR SPOC model with the crystal structure of PHF3 SPOC bound to RNA Pol II CTD di-heptapeptide. (**D**) Schematic of the HUSH complex with available structures overlaid.

In the AlphaFold model of the MPP8-TASOR complex, the TASOR subunit contains a SPOC domain with very high confidence scores (residues 352-502, pLDDT > 90), and a domain with a novel fold, DomI (residues 518-632, pLDDT > 70-90). DomI consists of four α-helices, with an extended 31-amino acid structured loop between the second and third helices. The first, second and fourth α-helices of DomI form a bundle with hydrophobic core consisting mostly of aromatic residues (**Fig. S3B**). These structural elements of DomI together occupy an extensive binding footprint on the PINIT-like domain of MPP8 (**Fig. 2B,C** and **Fig. S3A-C**). Comparison with the structure of yeast Siz1 in complex with PCNA, SUMO, and Ubc9 (Streich & Lima, 2016) shows that Siz1 uses a similar set of surfaces it its PINIT domain to interact with PCNA and Ubc9 as MPP8 uses in its PINIT-like domain to interact with TASOR DomI (**Fig. S3D**).

The SPOC domain is predicted to bind to the concave surface of the MPP8 ankyrin repeats. On the opposite side of the SPOC domain, residues 536-562 of MPP8 bind in an extended conformation the SPOC domain, extending the MPP8-TASOR interface (**Fig. 2B** and **Fig. S3A**). The 16-amino acid linker that connects the SPOC and DomI domains does not form any contacts with MPP8 and is predicted to be disordered (pLDDT = 30-50; **Fig. 2A**). MPP8 residues 547-562 were present in the crystallized MPP8 construct but were disordered in the electron density map. Together, the large total interface area of 4,374 Å^2^, the estimated solvation free energy gain, Δ*G*, of −54 kJ/mol, and the presence of 52 polar intermolecular contacts suggest that MPP8 and TASOR form a tight complex (**Fig. 2B-D** and **Fig. S3A-C**).

The AlphaFold model of the MPP8-TASOR complex is experimentally supported by our SEC data with different MPP8 and TASOR constructs (**Fig. S2B**), along with our previous coimmunoprecipitation and lentivirus reporter assays with different truncations of MPP8 and TASOR (Douse *et al*., 2020) (**Fig. 3**). In combination, these binding and functional assays identify the complex forming regions to MPP8 residues 547-860 and TASOR residues 354-633. Additionally, MPP8 residues 521-559, while not essential for TASOR binding, are required for HUSH repression activity (**Fig. 3**). The MPP8-TASOR AlphaFold model is consistent with these experimentally determined complex forming regions and functional boundaries (**Fig. 2A**). The only part of the predicted MPP8-TASOR interface that falls outside the experimentally determined complex forming regions is MPP8 residues 536-562 predicted to bind the TASOR SPOC domain (**Fig. 2B,C**). However, while these residues are dispensable for MPP8-TASOR binding, they are required for HUSH activity (**Fig. 3**).

**Fig. 3.**
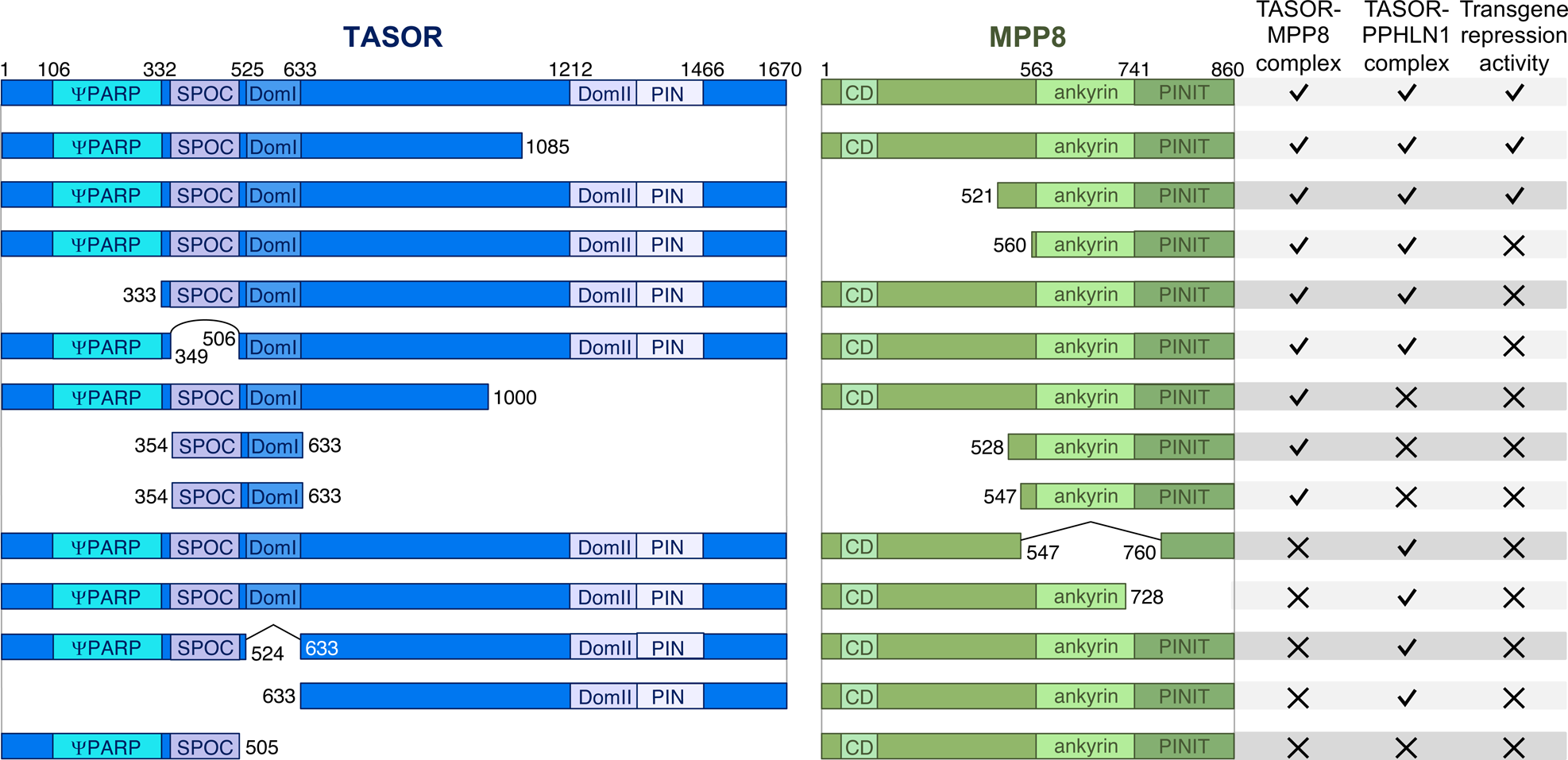
Effects of truncations in MPP8 and TASOR on HUSH complex formation and transgene repression. For complexes with TASOR residues 354-633, complex formation was assayed by size-exclusion chromatography in this study (**Fig. S2B**). For all other complexes, complex formation was assayed by co-immunoprecipitation in our previous study (Douse *et al*., 2020). Transgene repression activity was measured with an integrating lentivirus reporter in our previous study (Douse *et al*., 2020).

The most similar experimentally determined structures to the predicted TASOR SPOC domain are the SPOC domains from the *Arabidopsis* flowering regulator FPA, and human PHF3, with DALI Z-scores (Holm & Rosenstrom, 2010) of 12.2 and 11.8, respectively. Notably, the PHF3 SPOC domain binds RNA Pol II CTD di-heptapeptide phosphorylated on Ser2 (Y[pS]PTSPS-Y[pS]PTSPS) with a 1 µM dissociation constant. Binding of PHF3 SPOC to Ser2-phosphorylated Pol II CTD heptapeptide repeats, a hallmark of the elongating form of Pol II, has been proposed to promote Pol II elongation and condensation with mRNA processing factors, thereby contributing to the coordination of transcription elongation with mRNA decay (Appel *et al*, 2021). Superposition of the PHF3-Pol II CTD structure onto the predicted MPP8-TASOR structure shows that MPP8 residues 540-554 bind to the equivalent site on the TASOR SPOC domain as the Pol II CTD peptide on the PHF3 structure, such that binding of MPP8 and Pol II CTD heptapeptide to the TASOR SPOC domain would be mutually exclusive (**Fig. S3E,F**). The arginine and two lysine residues in the PHF3 SPOC domain that coordinate the phosphoserine moieties of the Pol II CTD are not conserved in the TASOR SPOC domain. Moreover, AlphaFold predictions with TASOR SPOC and Pol II CTD di-heptapeptides did not yield a consistent solution (**Fig. S3G**) and had very low confidence scores (pLDDT = 10-30). Based on these structural analyses, we conclude that it is unlikely that the TASOR SPOC domains binds to phosphorylated Pol II CTD heptapeptides like the PHF3 SPOC domain. Whether the TASOR SPOC domain recognizes another protein or peptide in related to transcription elongation or mRNA processing remains to be determined.

### The MPP8 CD binds H3K9me3-like peptides from H3K9 methyltransferases

The chromodomain (CD) of MPP8 binds H3K9me3 with relatively low affinity, with a dissociation constant (*K*_D_) in the micromolar range (Bua *et al*., 2009; Chang *et al*., 2011a; Quinn *et al*., 2010), and binding of this mark by MPP8 CD is not absolutely required for HUSH-dependent repression of lentiviruses in cell culture (Douse *et al*., 2020). The MPP8 CD binds human Dnmt3aK47me3 and the equivalent mark in mouse with even lower affinity (*K*_D_ = 12 µM) (Chang *et al*., 2011b), but whether the MPP8 CD binds directly to H3K9me3-like marks in other proteins remains unknown. To address this, we measured the binding affinities of MPP8 CD to the H3K9-like sequences in SETDB1, ATF7IP, G9a, and GLP. Our isothermal titration calorimetry (ITC) data show that MPP8 CD binds to the N-terminal tail of ATF7IP trimethylated at lysine 16 with a *K*_D_ of 193 ± 31 nM, and to residues 199-211 from GLP with lysine 205 trimethylated with a *K*_D_ of 337 ± 64 nM (**Fig. 4A**). These binding affinities are fourfold and twofold higher than for H3K9me3, which had a *K*_D_ of 769 ± 65 nM in ITC, consistent with previously reported values (Bua *et al*., 2009; Chang *et al*., 2011a; Quinn *et al*., 2010). MPP8 CD also bound to lysine-methylated peptides from G9a (trimethylated at K185) and SETDB1 (trimethylated at K1170) with *K*_D_ values of 1.31 and 1.55 µM, respectively (**Fig. 4A**). These affinities are slightly lower than H3K9me3 but slightly higher than H3K9me2 (*K*_D_ = 1.70 µM). Additionally, H3K27me3 and SETDB1 K1178me3 peptides showed weak binding (*K*_D_ > 50 µM). There was no detectable binding of MPP8 CD to any of the peptides listed above in the absence of lysine methylation (**Fig. 4A,B** and **Fig. S4**). In summary, our ITC data shows that the MPP8 CD binds to lysine-trimethylated peptides in all four components of the SETDB1/ATF7IP and G9a/GLP H3K9 methyltransferase complexes, with binding affinities higher than for H3K9me3 in the case of the ATF7IP and GLP peptides.

**Fig. 4.**
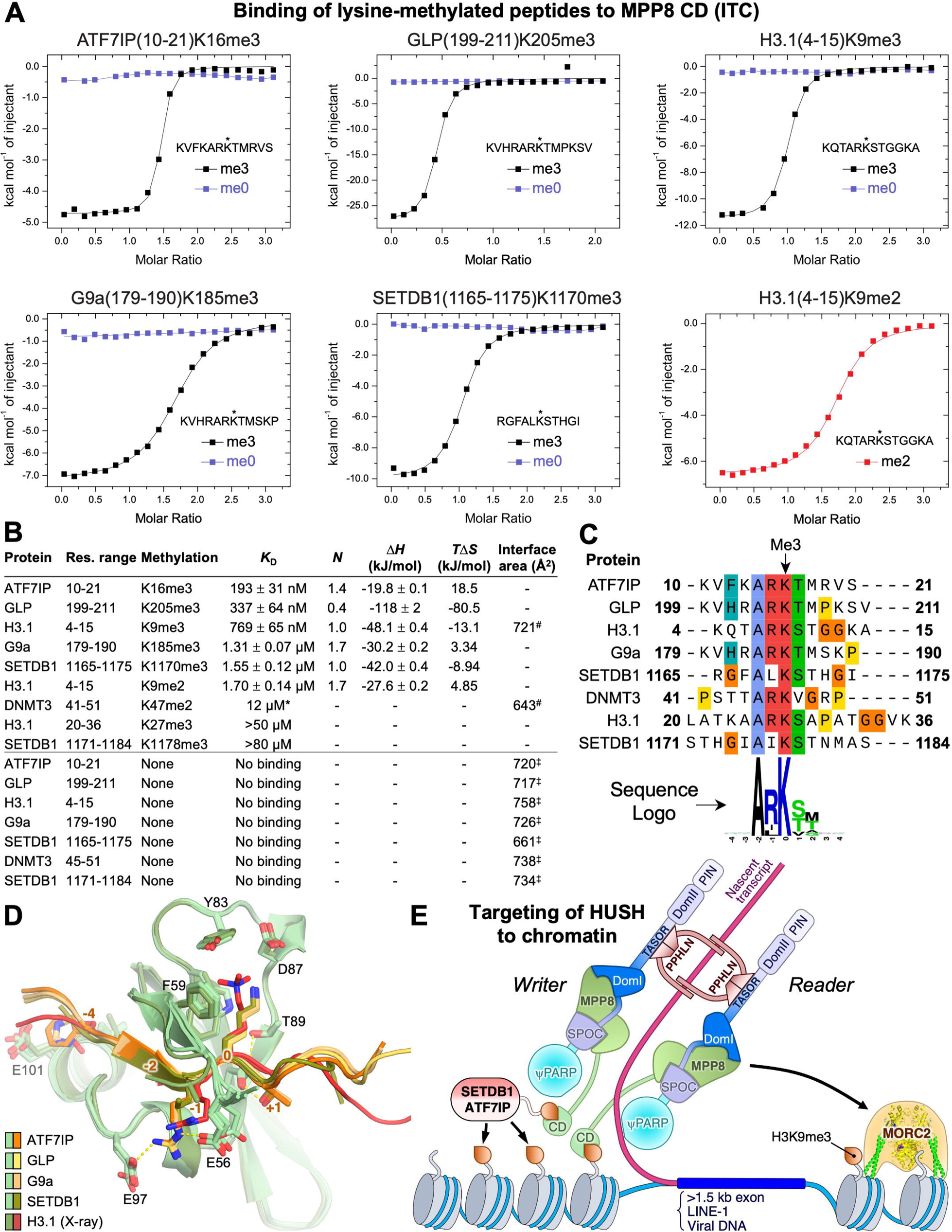
Binding of the MPP8 CD to H3K9 and H3K9-like peptides. (**A**) Isothermal titration calorimetry (ITC) thermograms from titration into a solution of MPP8 CD of six peptides, either unmethylated (blue) or trimethylated at the lysine residue marked with an asterisk (black). See **Fig. S4** for the complete set of ITC isotherms and thermograms and data for additional peptides. (**B**) Dissociation constants (*K*_D_), binding stoichiometry (*N*) and thermodynamic parameters derived from ITC data. The listed interface areas are for crystal structures (“#”) of MPP8 CD bound to H3K9me3 (PDB 3QO2) (Chang *et al*., 2011a) and hDnmt3aK47me2 (PDB 3SVM) (Chang *et al*., 2011b); and AlphaFold models (“‡”) of MPP8 CD bound to the indicated H3K9-like sequences. (**C**) Multiple sequence alignment of H3K9 and the H3K9-like sequences from the ITC measurements. UniProt accession numbers: ATF7IP, Q6VMQ6; GLP, Q9H9B1; H3.1, P68431; G9a, Q96KQ7; SETDB1, Q15047; DNMT3a, Q9Y6K1. Lower panel: sequence logo generated with WebLogo (Crooks *et al*, 2004). (**D**) Superposition of MPP8 CD-peptide AlphaFold models on the crystal structure with H3K9me3 bound. Numbers in orange show the residue position relative to the methyl-lysine. See **Fig. S5C** for a closeup of the CD-peptide interactions. (**E**) Model for targeting of the HUSH complex with multivalent engagement to actively transcribed retrotransposons.

Having identified nine lysine-methylated peptides with measurable binding to the MPP8 CD (**Fig. 4B**) we generated atomic models of the MPP8-peptide complexes with AlphaFold-Multimer. Although lysine methylation could not be modeled in AlphaFold, we expected the modeling to provide insights on how the amino acid sequence flanking the lysine methylation contributes to MPP8 binding. The MPP8 CD was reported to form homodimers in SEC and in crystals in complex with an H3K9me3 peptide (Chang *et al*., 2011a). To obtain a more accurate measurement of the molecular weight of the MPP8 CD in solution independent of its shape, we measured multiangle light scattering during SEC (SEC-MALS). We obtained the same SEC profile as previously reported (Chang *et al*., 2011a), but the MALS data unambiguously showed that the MPP8 CD was a monomer rather than a dimer (**Fig. S5A**). Consistent with the MPP8 CD being monomeric in solution, analysis with the PISA (39) showed that the crystallographic MPP8 CD dimer interface area (534 Å) and estimated solvation free energy gain (ΔG = −3.6 kcal/mol) were both smaller than would be expected for the interface to be stable in solution at physiological concentrations of MPP8. Moreover, AlphaFold-Multimer predictions with two copies of the MPP8 CD, without H3K9-like peptide or use of PDB templates, failed to consistently predict the crystallographic MPP8 dimer. We therefore specified a 1:1 MPP8:peptide stoichiometry for AlphaFold modeling. The resulting MPP8-peptide complexes had high confidence scores, with an average pLDDT of 73 for the peptides and 87-91 for the MPP8 CD (**Fig. S5B**). For each complex, the peptide and MPP8 CD in the top five predicted models generated without PDB templates had the same overall structure as the previously determined crystal structure of the MPP8 CD bound to an H3K9me3 peptide (Chang *et al*., 2011a) (**Fig. 4D**). In all the structures, the lysine at the methylation site (position 0) is bound in a cage formed by an aspartate and three aromatic residues from MPP8 (**Fig. 4C-D** and **Fig. S5C**). All peptides bound by MPP8 contain an alanine a position −2 (**Fig. 4C**). The side chain of this alanine packs onto a hydrophobic surface of the CD, forming van der Waals contacts. There is a strong preference of an arginine at position −1, with only the two SETDB1 peptides lacking an arginine at this position. In the AlphaFold models, the side chain of the arginine at position −1 is predicted to form a salt bridge with Glu56 of MPP8, although we note that the Glu56 side chain was disordered in the MPP8-H3K9me3 crystal structure (**Fig. 4D** and **Fig. S5C**). Position +1 is a serine or threonine in all peptides except the Dnmt3a peptide, which has a valine at this position. The hydroxyl of this serine or threonine contributes to MPP8 binding by forming a hydrogen bond with Glu91 of MPP8. The peptides with highest affinities for MPP8, those from ATF7IP, GLP, and G9a, also contain an aromatic residue (histidine or phenylalanine) at position −4, which is predicted to form hydrophobic contacts with the C-terminal helix of the MPP8 CD. In contrast, the H3K9me3 peptide has a glutamine at position −4. AlphaFold predictions with scrambled (randomized) amino acid sequences of the ATF7IP, GLP, G9a and SETDB1 peptides resulted more variable peptide conformations with non-lysine residues in the trimethyl-lysine binding cage (**Fig. S5D**). Additionally, in most of these predictions the scrambled peptides were bound in the opposite direction to the experimental and predicted structures with native peptide sequences. Together, our structural modeling suggests that the key determinants for recognition of a methylated lysine by MPP8 are a strict requirement for an alanine at position −2, strong preferences for arginine at position −1 and serine or threonine at position +1, and a possible preference for an aromatic residue at position −4.

## Discussion

The crystal structure of the MPP8 CTD and the AlphaFold models of the MPP8-TASOR complex complete our picture of the of the folded domains essential for HUSH repression of integrated transgenes. This leaves the DomII and PIN domains of TASOR (residues 1212-1466), which are not required for transgene repression, as the only domains in the HUSH complex that are predicted to be folded but structurally unstudied. Our model of the MPP8-TASOR complex identifies an extensive molecular interaction interface between the MPP8 CTD and the SPOC and DomI domains of TASOR. The model reveals structural homologies between the MPP8 CTD and the substrate recognition domains of Siz/PIAS-family SUMO E3 ligases, and between the SPOC domains of TASOR and PHF3, a protein that contributes to the linkage of transcription elongation with mRNA decay. The TASOR DomI domain is predicted to form a novel fold that is structurally dependent on the MPP8 CTD. A long loop in DomI that binds along a large area of the PINIT-like domain of MPP8 contains a lysine (Lys586) reported to be SUMOylated (Hendriks *et al*., 2017). Lys586 is highly solvent accessible in the predicted structure (**Fig. 2C** and **Fig. S3B**), suggesting that a function of the MPP8 PINIT-like domain could be to present TASOR K586 as a substrate for a SUMO E3 ligase. This raises the question of whether SUMOylation of TASOR Lys586 may regulate or license transcriptional repression by the HUSH complex.

Our ITC measurements show that the CD of MPP8 binds to lysine-trimethylated H3K9-like peptides in all four components of two key H3K9 methyltransferases, SETDB1, ATF7IP, G9a, and GLP, with slightly higher affinity than H3K9me3. AlphaFold modeling suggests that the MPP8 CD preferentially recognizes the lysine trimethylated peptides with the sequence ARK[S/T], with a possible preference for an aromatic residue at position −4. A reported phosphorylation site in the MPP8 CD, at Thr89 (Ochoa *et al*, 2020), maps to the vicinity of the methyl-lysine binding site (**Fig. S5C**). The structures of the CD-peptide complexes suggest that a phosphothreonine adduct at this site would be compatible with H3K9-like peptide binding but is likely to alter the binding affinity and potentially also the methylation specificity of MPP8. The interaction of MPP8 with trimethylated peptides from H3K9 methyltransferases suggests that the MPP8 CD can recruit SETDB/ATF7IP to target chromatin loci in addition to sensing (reading) H3K9me3. The H3K9-like peptides from ATF7IP, G9a and GLP are methylated by the G9a/GLP complex, and these methylated peptides can then bind the CD of HP1 (Chang *et al*., 2011b; Chin *et al*., 2007; Kokura *et al*., 2010; Sampath *et al*., 2007; Tsusaka *et al*., 2018). Thus, recruitment of H3K9 methyltransferases by MPP8 may promote recruitment of HP1 and heterochromatinization through amplification of lysine trimethylation not only on H3K9 but also on H3K9-like peptides in the H3K9methyltransferases themselves. However, binding of the MPP8 CD to H3K9me3 is not absolutely required for targeting HUSH to its chromatin loci, as HUSH can bind chromatin in SETDB1 knockout cells, through binding of Periphilin to nascent transcripts (Seczynska *et al*., 2022).

Our biochemical, structural, and modeling data have major implications for the assembly and oligomeric state of the HUSH complex. We found that MPP8 and TASOR form a heterodimer. We showed previously that in the core Periphilin-TASOR complex two molecules of Periphilin bind to a single copy of TASOR (Prigozhin *et al*., 2020). Consequently, the HUSH complex has a stoichiometry of 1:1:2 MPP8:TASOR:Periphilin, or a multiple thereof. Self-aggregation of the N-terminal domain of Periphilin will draw in additional copies of Periphilin and hence incorporate Periphilin-bound nascent transcripts into higher-order molecular condensates (Prigozhin *et al*., 2020). These additional Periphilin molecules could recruit additional copies of MPP8 and TASOR, which would allow simultaneous engagement of HUSH with H3K9me3 and SETDB1/ATF7IP via MPP8, with one copy of MPP8 bound to (reading) an H3K9me3 peptide and the other copy of MPP8 bound to SETDB1/ATF7IP, which could then deposit (write) new H3K9me3 marks. Alternatively, this higher-order assembly may serve to increase the stability and local concentration of HUSH subunits and effectors by providing multivalent interactions between the proteins in the assembly, and with nucleosomes and nascent transcripts at target loci. Stabilization of HUSH-methyltransferase complexes through multivalency, or avidity, could explain why mutation of the lysine in the H3K9-like peptide of ATF7IP caused a loss of MPP8 chromatin binding and delayed HUSH-dependent repression of a reporter transgene in mouse embryonic stem cells (Tchasovnikarova *et al*., 2015; Tsusaka *et al*., 2018). Similarly, a conditional knockout of SETDB1 led to loss of MPP8 recruitment to HUSH target loci along with a reduction in H3K9me3 (Muller *et al*., 2021). Therefore, we propose that multivalent binding of the HUSH complex via MPP8 to SETDB1/ATF7IP contributes to chromatin binding of the complex (**Fig. 4E**). Complicating this interpretation, however, there is evidence that MPP8 can bind chromatin outside of the HUSH complex, as a subset of MPP8 chromatin binding sites lack TASOR and Periphilin (Spencley *et al*., 2023). Consistent with this, MPP8 recruits the NEXT complex to chromatin via an interaction with ZCCHC8, leading to exosome-dependent decay of shorter (<1 kb) non-polyadenylated transcripts, whereas HUSH represses polyadenylated transcripts (Garland *et al*., 2022).

The data presented here allows us to propose an updated model for transcriptional repression by the HUSH complex (**Fig. 4E**). The N-terminal domain of Periphilin self-aggregates and binds nascent transcripts with intronless, adenine-rich gene bodies longer than 1.5 kb, resulting in large ribonucleoparticles or condensates containing multiple copies of Periphilin bound to each transcript. The pseudo-PARP domain of TASOR may also contribute to transcript binding (Douse *et al*., 2020). Transcripts within these ribonucleoparticles would be less accessible to mRNA processing and export machinery, thereby repressing expression. Since ribonucleoparticles will contain many copies of Periphilin, multiple copies of TASOR and then MPP8 are recruited. Some of the MPP8 CDs will bind H3K9me3, if present, and other MPP8 subunits will recruit SETDB1/ATF7IP and possibly G9a/GLP complexes via interactions between the MPP8 CD and methylated H3K9-like sequences in the methyltransferase components. The recruited SETDB1 deposits new H3K9me3 marks providing a read/write amplification mechanism. MORC2 is subsequently recruited to drive chromatin compaction and remodeling. How MORC2 is recruited remains unclear. Many other questions remain, such as for example what the HUSH-independent function of MPP8 may be, how domains other than the MPP8 CD contribute to recruitment of HUSH to retrotransposons, the roles of the TASOR pseudo-PARP and DomII/PIN domains, and how the requirements for the establishment of transcriptional repression by the HUSH complex may differ from the requirements to maintain repression, once established. Further studies are warranted to address these questions and test the model proposed above.

The HUSH complex fulfills a function vital to the cell in protecting the genome and repressing expression of integrated retroelements of endogenous or viral origin. Our work identifies novel structural and biochemical properties of MPP8 that are necessary for HUSH complex assembly and transcriptional repression. In the context of previous studies, the data presented here allows us to update the working model of HUSH-dependent transcriptional repression to include multivalent binding to H3K9me3 and H3K9-like sequences in methyltransferases resulting in spreading of the H3K9me3 mark through a read/write mechanism. HUSH inhibitors could be useful to stimulate autoinflammation in cancer immunotherapy or help bring retroviruses out of latency so they can be treated with antiretrovirals. Targeting the MPP8 CD may be useful in applications that require fractional inhibition of HUSH activity to limit toxicity. A family of peptidomimetic MPP8 CD ligands has been identified, providing proof of principle that MPP8 CD-specific molecules can be developed (Waybright *et al*, 2021). Our structural data provide a foundation for further design and optimization of HUSH inhibitors targeting MPP8.

## Methods

### Protein expression and purification

Synthetic genes codon-optimized for *Escherichia coli* were cloned into the pET15b vector (Novagen) with an N-terminal hexahistidine (His_6_)-tag. The MPP8 CTD (residues 547-860, UniProt Q99549) was cloned in frame with the vector’s His_6_ tag and thrombin cleavage site. The MPP8 CD (residues 55-116) was cloned with a tobacco etch virus (TEV) protease cleavage site (ENLYFQG) replacing the thrombin cleavage site. These constructs were transformed into *E. coli* BL21 (DE3) cells (New England BioLabs).

For expression and purification of MPP8 CTD, overnight *E. coli* cultures were diluted 1:200 into 2 liters of autoinduction media (25 mM Na_2_HPO_4_, 25 mM KH_2_PO_4_, 50 mM NH_4_Cl, 5 mM Na_2_SO_4_, 2 mM MgSO_4_, 0.2 x trace metals mix, 0.5% glycerol, 0.05% glucose, 0.2% α-lactose, 0.1% aspartate, 0.2 mg/ml of 18 amino acids (no C or Y), 1 µM vitamin B12, 0.1 mg/ml ampicillin) and grown at 37°C for 3 h and then 18°C for 26 h. Cells were harvested and resuspended in 50 ml Lysis-Wash buffer (20 mM Tris-HCl pH 7.4, 0.15 M NaCl, 1 mM DTT (1,4-dithiothreitol), 20 mM imidazole). Cells were lysed by sonication and centrifuged at 16,000 rpm for 20 min. The supernatant was filtered through a 0.45 µm syringe filter and applied to a 5 ml Ni-NTA agarose column (Qiagen) pre-equilibrated in Lysis-Wash buffer. The column was washed with 50 ml Lysis-Wash buffer. MPP8 CTD was eluted with 25 ml of elution buffer (20 mM Tris-HCl pH 7.4, 0.15 M NaCl, 1 mM DTT, 0.4 M imidazole). To remove the His_6_ tag, 0.1 ml of thrombin (Cytiva) was added to the eluate and the mixture was incubated overnight at room temperature with concomitant dialysis against 1 liter of dialysis buffer (20 mM Tris-HCl pH 7.4, 20 mM NaCl, 1 mM DTT, 10% glycerol). The solution was applied to a column of 5 ml Ni-NTA agarose (Qiagen) equilibrated in Lysis-Wash buffer and the flow-through collected. The sample was then applied to a Resource Q anion exchange column (Cytiva) equilibrated in low-salt buffer (20 mM Tris-HCl pH 8, 20 mM NaCl, 1 mM DTT, 10% glycerol). The protein was eluted with a linear gradient of high-salt buffer (10 mM HEPES pH 7.4, 1 M NaCl, 0.5 mM Tris(2-carboxyethyl)phosphine (TCEP)). Fractions containing MPP8 CTD were pooled, concentrated, and applied to a Superdex 200 10/300 size-exclusion chromatography (SEC) column (Cytiva) equilibrated in SEC buffer (10 mM HEPES pH 7.8, 0.15 M NaCl, 0.5 mM TCEP).

To co-express TASOR and MPP8 to assay for complex formation, we transformed *E. coli* BL21(DE3) cells with a pRSF vector (Novagen) encoding TASOR (residues 354-633) and the pET15b vector encoding different MPP8 CTD fragments. Cells were cooled to 18°C at an optical density (OD_600 nm_) of 0.6 and induced with 0.1 mM IPTG (isopropyl β-D-1-thiogalactopyranoside) at OD_600 nm_ = 0.8. After overnight incubation, cells were harvested and resuspended in 20 mM HEPES pH 7, 0.5 M NaCl, 0.5 mM TCEP, 20 mM imidazole. Cells were lysed and purified by nickel-affinity chromatography as above. Eluted TASOR-MPP8 complexes were analyzed by SEC on a Superdex 200 10/300 column in 20 mM HEPES pH 7.5, 0.5 M NaCl, 0.5 mM TCEP.

To express and purify MPP8 CD, overnight *E. coli* BL21 (DE3) cultures were diluted 1:200 into 1.6 liters of LB with 0.1 mg ml^−1^ ampicillin, grown at 37°C to OD_600_ 0.6, induced with 0.1 mM IPTG, and incubated at 37°C for 4-6 h. Cells were harvested and resuspended in twice the cell pellet volume of lysis buffer (50 mM potassium phosphate pH 8.0, 0.5 M NaCl, 20 mM Imidazole, cOmplete protease inhibitor cocktail (Roche)). Cells were lysed by sonication and centrifuged at 35,000 rpm for 1 h. The supernatant was mixed with 2 ml Ni-NTA agarose (Qiagen) pre-equilibrated in lysis buffer. The mixture was incubated for 1 h, washed with wash buffer (50 mM potassium phosphate pH 8.0, 0.5 M NaCl, 20 mM imidazole), and bound protein was eluted with wash buffer supplemented with 0.3 M imidazole. The buffer was exchanged by serial dilution and concentration to Mono Q buffer (25 mM Tris pH 7.4, 50 mM NaCl, 0.5 mM TCEP). The sample was applied to a Mono Q 5/50 GL column (Cytiva) equilibrated in Mono Q buffer. Bound protein was eluted with a linear gradient of high-salt Q buffer (25 mM Tris pH 7.4, 1 M NaCl, 0.5 mM TCEP). Fractions containing MPP8 CD were pooled, concentrated, and applied to a Superdex 75 10/300 SEC column (Cytiva) equilibrated in SEC75 buffer (50 mM HEPES pH 7.5, 0.15 M NaCl, 0.5 mM TCEP).

### Crystallographic structure determination

Crystals were grown at 18°C by sitting drop vapor diffusion. Purified MPP8 CTD was concentrated to 4.5 mg/ml using a 10-kDa cutoff centrifugal filtration device (Merck Millipore) and mixed with an equal volume (up to 0.5 µl) of reservoir solution. Prism- and needle-shaped crystals were seen in 13 conditions from the Morpheus II screen (Molecular Dimensions), each containing 0.5 mM Oxometalates Mix and either 10% w/v PEG 8000 with 20% v/v 1,5-pentanediol or 12.5% w/v PEG 4000 with 10% w/v 1,2,6-hexanetriol. Crystals suitable for X-ray diffraction experiments were obtained with the following Morpheus II reagents: Precipitant Mix 6 (1:2 dilution), Buffer System 6 pH 8.5 (1:10 dilution), and Oxometalates Mix (1:10 dilution). The final precipitant solution contained 12.5% (w/v) PEG 4000, 20% (v/v) 1,2,6-hexanetriol, 0.5 mM of each Oxometalate, 0.1 M GlyGly/AMPD pH 8.5. Microseeding was used to obtain larger crystals; the seed stock was made by shearing small crystal by repeated pipetting and performing serial dilutions of the sheared crystal fragments to use as seed stocks. Incubation of crystals in mother liquor supplemented with 10-20% PEG200 for 30 min prior to freezing in liquid nitrogen slightly improved the X-ray diffraction limit to 3.0 Å resolution.

X-ray diffraction data were collected at 100 K at Diamond Light Source (DLS) beamline I04-1 and processed with Xia2 (Dials, Aimless) (Winter, 2010). An atomic model predicted by AlphaFold2 as implemented by ColabFold (Mirdita *et al*., 2022) was used as a molecular replacement search model in Phenix v.1.20 (Adams *et al*, 2010). Eight copies of the model were placed in the asymmetric unit (chains A-H). The atomic coordinates were edited with Coot v.0.9.6 (Emsley & Cowtan, 2004) and iteratively refined with Phenix (Adams *et al*., 2010). The final refined atomic coordinates of the crystal structure spanned residues 563-860 (residues 547-562 were disordered). See **Table 1** for crystallographic data collection, refinement, and validation statistics.

### Isothermal titration calorimetry (ITC)

H3K9 and H3K9-like peptides, unmethylated or lysine-trimethylated, were chemically synthesized (Bio Basic Inc / NBS Biologicals Ltd). Binding of MPP8 CD to H3K9 and H3K9-like peptides was analyzed in 50 mM HEPES pH 7.4, 0.1 M NaCl, 0.5 mM TCEP at 25°C (298 K), with an AutoiTC200 calorimeter (MicroCal). The sample cell was loaded with 0.37 ml of 50 µM MPP8 CD and the titrant syringe with 750 µM peptide. 20 serial injections of 6 μl peptide solution were performed at 3 min intervals. The stirring speed was 1,000 rpm and the reference power was maintained at 6.06 μcal s^−1^. The net heat absorption or release associated with each injection was calculated by subtracting the heat associated with the injection of peptide solution to buffer. Thermodynamic parameters were extracted from a curve fit to the data to a single-site model with Origin 7.0 (MicroCal).

### Size-exclusion chromatography and multiangle scattering (SEC-MALS)

100 µl samples containing 4 mg ml^−1^ MPP8 CD were analyzed by size-exclusion chromatography (SEC) at 293 K on a Superdex 75 (10/300) column (Cytiva) in 50 mM HEPES pH 7.5, 0.15 M NaCl, 0.5 mM TCEP with a flow rate of 0.5 ml min^−1^ on an Agilent 1200 series liquid chromatography system. The SEC system was coupled to a Wyatt Heleos II 18-angle light scattering instrument coupled to a Wyatt Optilab T-rEX online refractive index detector. Protein concentration was determined from the excess differential refractive index based on 0.185 dRI for 1 g ml^−1^ protein (dn/dc = 0.185 ml g^−1^). The measured protein concentration and scattering intensity were used to calculate molecular mass from the intercept of a Debye plot using Zimm’s model as implemented in the Wyatt Astra software. Bovine serum albumen (100 µl at 2 mg ml^−1^) was run as a standard for mass determination, and to determine interdetector delay volumes and band broadening parameters.

### AlphaFold2 modeling

Atomic models of MPP8 CTD in complex with TASOR SPOC-DomI were predicted with a local installation of AlphaFold2-v2.3.1 (Evans *et al*., 2021; Jumper *et al*., 2021) using multimer mode as implemented by ColabFold v1.5.2 (Mirdita *et al*., 2022) using MMseqs2 for multiple sequence alignment (MSA). The amino acid sequences of human MPP8 residues 535-860 and TASOR residues 352-632 (UniProt Q9UK61-1) were used as the input. Polypeptide geometry in the models was regularized by post-prediction relaxation using the Amber force field. The maximum number of recycles was set to 20. No MSA cutoff was used. The sequence pairing mode was set to pair sequences from the same species and unpaired MSA. The multimer model was set to multimer-v3. Use of available PDB template files had no significant effect on atomic model prediction. Five atomic models were output from each AlphaFold run. Protein interfaces were analyzed with the Protein interfaces, surfaces, and assemblies (PISA) service at the European Bioinformatics Institute [http://www.ebi.ac.uk/pdbe/prot_int/pistart.html] (Krissinel & Henrick, 2007).

Atomic models of MPP8 CD in complex with H3K9 and H3K9-like peptides were generated as described above except that the input sequences consisted of human MPP8 CD (residues 55-116) and one of the following: ATF7IP residues 10-21 (UniProt Q6VMQ6), GLP residues 199-211 (UniProt Q9H9B1), histone 3.1 residues 4-15 (UniProt P68431), G9a residues 179-190 (UniProt Q96KQ7), SETDB1 residues 1165-1175 (UniProt Q155047), and DNMT3a residues 45-51 (UniProt Q9Y6K1). Use of available PDB template files slightly improved the confidence scores of the models.

The top-ranking atomic coordinates generated by AlphaFold-Multimer for the MPP8-TASOR complex and for the MPP8-peptide complexes shown in this study are available in ModelArchive with accession codes ma-t4h1j, ma-awwwj, ma-jz826, ma-zbsvq, ma-mlhvc, and ma-58pyq.

## Acknowledgments

We thank Fabrice Gorrec (MRC-LMB Crystallization Facility) for advice on protein crystallization, and Dom Bellini at the MRC-LMB X-ray Crystallography Facility for advice in crystallographic data collection. We thank Chris Batters (MRC-LMB Biophysics Facility) for advice and training in SEC-MALS. We are grateful to Paul Lehner (University of Cambridge Department of Medicine) for helpful comments on the manuscript. Crystallographic data were collected on beamline I04-1 at Diamond Light Source (DLS). For the purpose of open access, the University of Cambridge has applied a CC BY public copyright licence to any Author Accepted Manuscript version arising. This work was supported by the Wellcome Trust [101908/Z/13/Z to Y.M.; 217191/Z/19/Z to Y.M.]. Access to Diamond Light Source was supported by the Wellcome Trust; the Medical Research Council; and UKRI [proposal number MX21426].

## Author Contributions

Conceptualization, D.M.P., S.O. and Y.M.; Methodology, N.N., D.M.P., S.O. and Y.M.; Investigation, N.N., D.M.P, S.O. and Y.M.; Validation – crystal structure, Y.M., Y.M.; Writing, Y.M.; Visualization, N.N. and Y.M.; Supervision, Y.M.; Project Administration, Y.M.; Funding Acquisition, Y.M.

## Data availability

The structure factors and atomic coordinates for the MPP8 CTD are available in the Protein Data Bank with code PDB: 8QFB, at https://dx.doi.org/10.2210/pdb8QFB/pdb. The original experimental X-ray diffraction images are available in the SBGrid Data Bank (data.SBGrid.org) with Data ID 1043, at https://dx.doi.org/10.15785/SBGRID/1043. The AlphaFold models are available in modelarchive.org at https://dx.doi.org/10.5452/ma-t4h1j, https://dx.doi.org/10.5452/ma-awwwj, https://dx.doi.org/10.5452/ma-jz826, https://dx.doi.org/10.5452/ma-zbsvq, https://dx.doi.org/10.5452/ma-mlhvc, and https://dx.doi.org/10.5452/ma-58pyq. Other data underlying this article are available in the article and Expanded View content.

## Supplementary Figure Legends

**Fig. S1.** Crystal structure of the MPP8 C-terminal domain (CTD). (**A**) Representative samples of the 2F_o_ – F_c_ electron density map of the crystal structure of MPP8 CTD. The eighth chain in the asymmetric unit (chain G) is shown. An isomesh contour level of 1.0 σ was used in PyMol (Schrödinger, LLC). (**B**) One of four disulfide linked CTD homodimers in the crystallographic asymmetric unit. Inset, closeup of the dimer interface.

**Fig. S2.** Oligomerization states of the MPP8 C-terminal domain (CTD) and MPP8-TASOR complexes. (**A**) Size-exclusion chromatography (SEC) of the MPP8 CTD. The peaks at 14 ml and 16 ml elution volumes both contain pure MPP8 CTD suggesting the two peaks correspond to dimer and monomer, respectively. (**B**) SEC of minimal TASOR-MPP8 complexes.

**Fig. S3.** Detailed properties of the AlphaFold-Multimer model of the MPP8-TASOR complex. (**A**) Predicted interactions between MPP8 and the SPOC domain of TASOR. (**B**) Predicted interaction between MPP8 and TASOR DomI. (**C**) Superposition of the five highest scoring AlphaFold models of the MPP8-TASOR complex. Flanking sequences were included to highlight the boundaries of the folded domains (marked with arrows). (**D**) Crystal structure (PDB 5JNE) of yeast Siz1 in complex with its substrate PCNA, SUMO ortholog SMT3, and SUMO E2 ligase Ubc9 (Streich & Lima, 2016). (**E**) Crystal structure (PDB 6IC8) of the PHF3 SPOC domain (light purple) bound to Ser2-phosphorylated RNA Pol II CTD di-heptapeptide (dark purple) (Appel et al, 2021). The phosphoserine side chains are shown in stick representation. (**F**) AlphaFold model of the TASOR SPOC domain (blue) in complex with MPP8 (green). (**G**) Superposition of the five highest scoring AlphaFold models of the TASOR SPOC domain (blue) in complex RNA Pol II CTD di-heptapeptide.

**Fig. S4.** ITC isotherms and thermograms for titration of H3K9 and H3K9-like peptides, lysine-methylated or unmethylated, into a solution of the MPP8 chromodomain (CD).

**Fig. S5.** Oligomerization state of MPP8 CD and structural modeling in complex with H3K9-like peptides with AlphaFold-Multimer. (**A**) Size-exclusion chromatography with multiangle light scattering (SEC-MALS) of MPP8 CD. dRI, relative refractive index. MW, molar mass. (**B**) AlphaFold predictions generated without using template atomic coordinates from the Protein Data Bank. Upper panels: atomic models of the chromodomain-peptide (CD-P) complexes colored by model confidence (pLDDT). Lower panels: predicted aligned error plots. (**C**) Closeup of the interactions between MPP8 CD and H3K9me3 or H3K9-like peptides. (**D**) Atomic models and predicted aligned error plots from AlphaFold-Multimer runs using a scrambled (randomized) amino acid sequence of each peptide in (B). The terminal residues of each peptide are numbered by distance from the residue bound to the trimethyl-lysine binding cage. This residue is underlined each peptide sequence.

