## Supplementary figures and images for "Structure and methyl-lysine binding selectivity of the HUSH complex subunit MPP8"

### Supplemental Figures

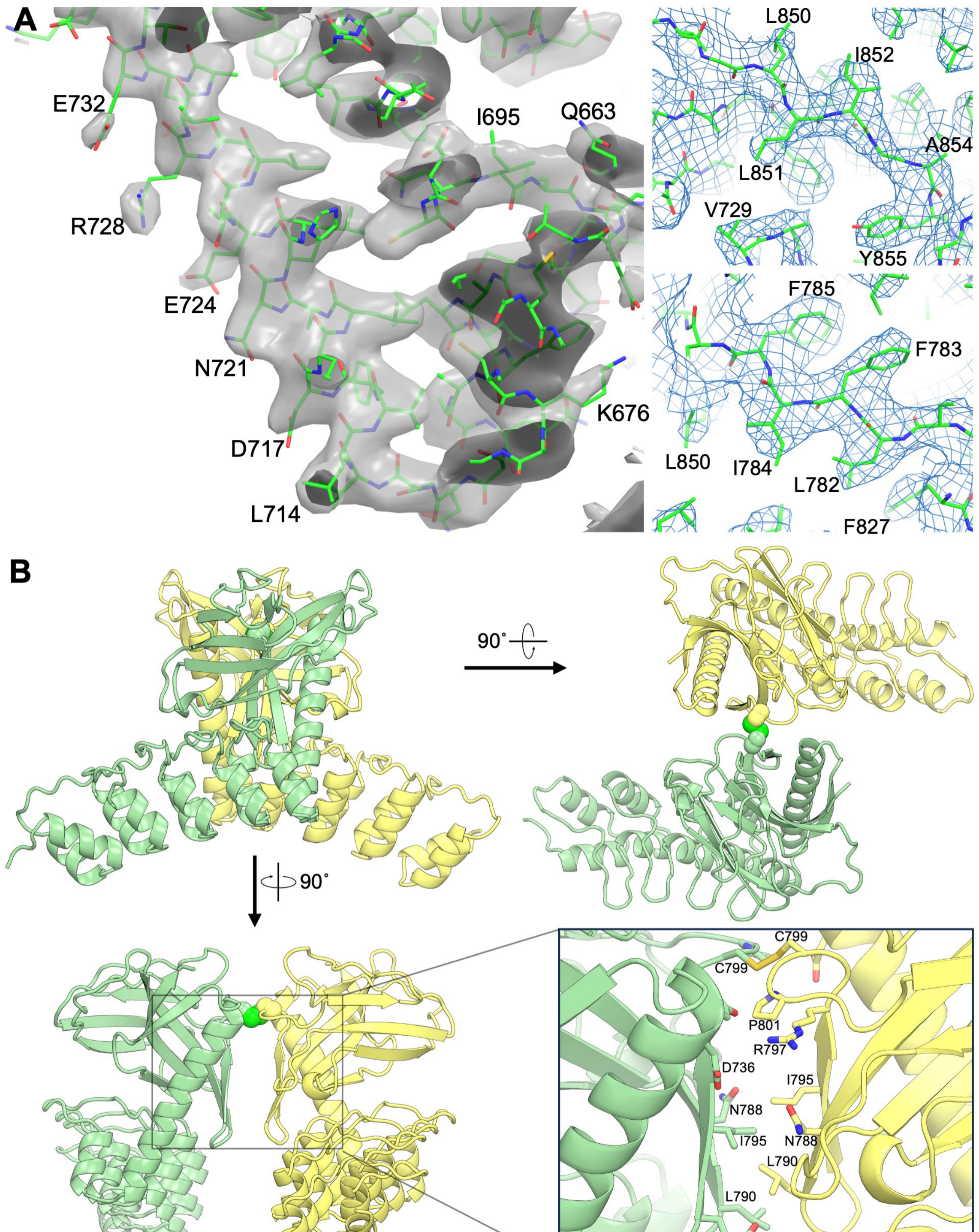


**Protein interface statistics**

| Area | $\Delta^iG$ | $\Delta^iG$ P-value | Polar contacts |
|---|---|---|---|
| 377 Å$^2$ | -10.8 kJ/mol | 0.014 | 1 |

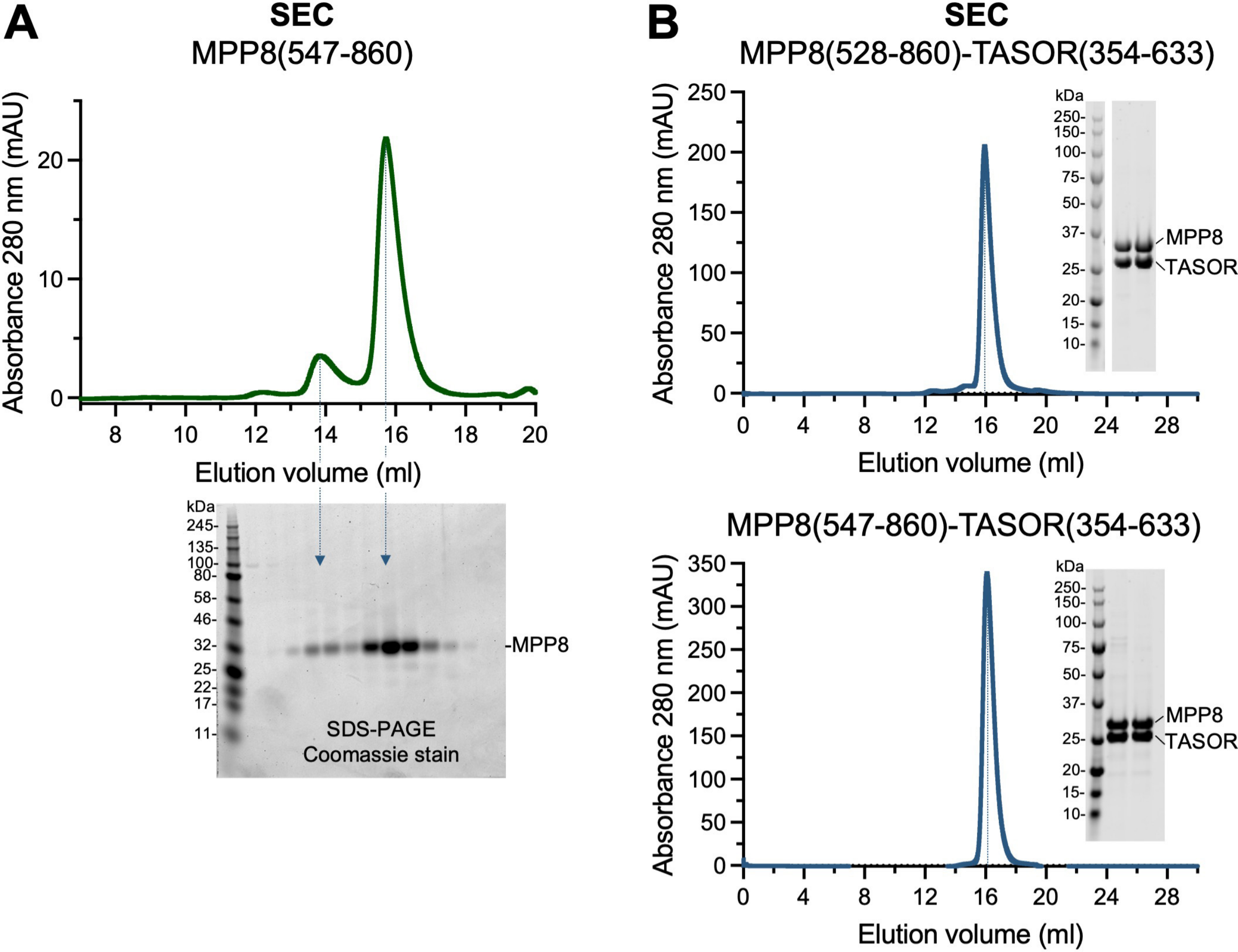

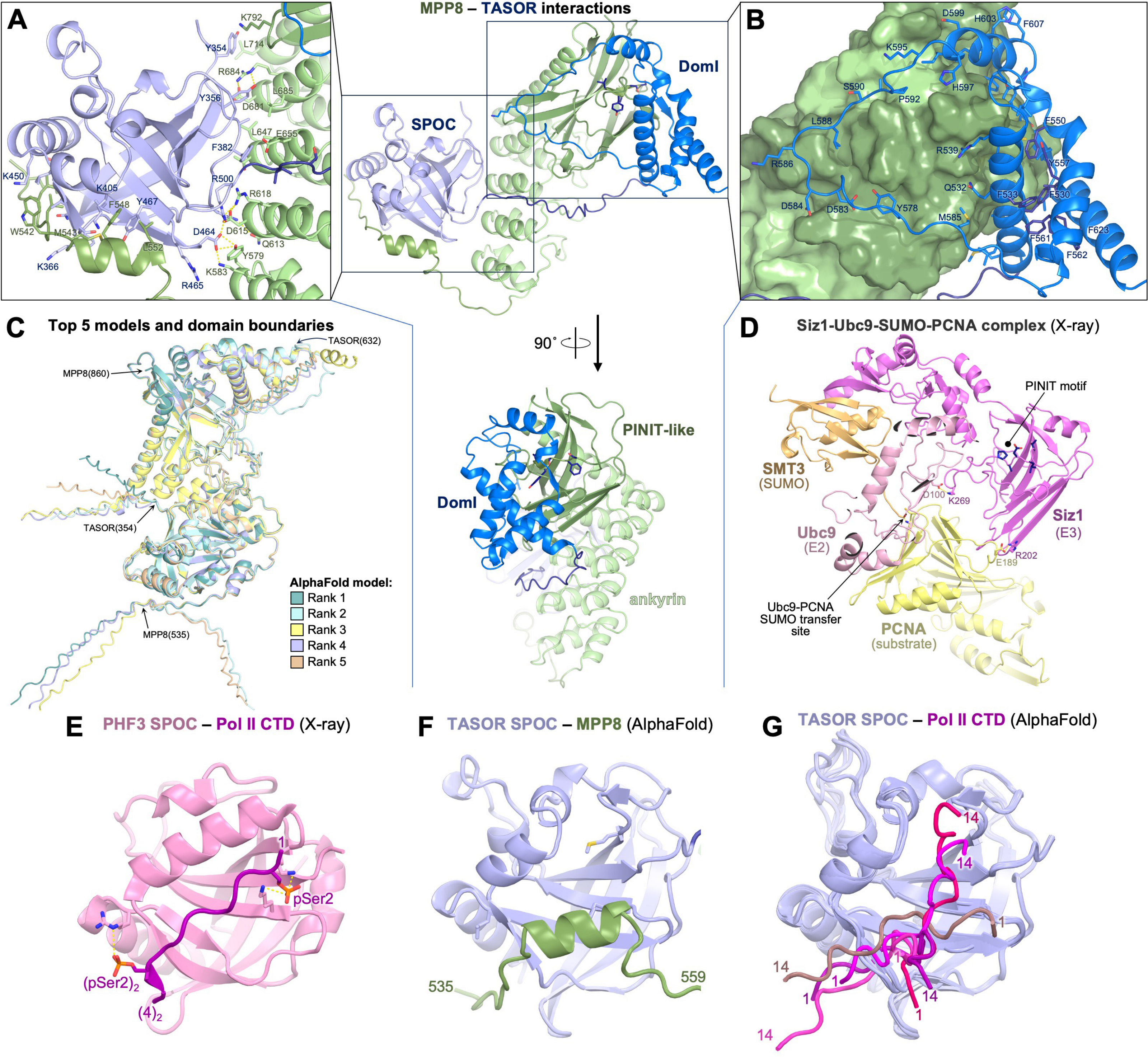

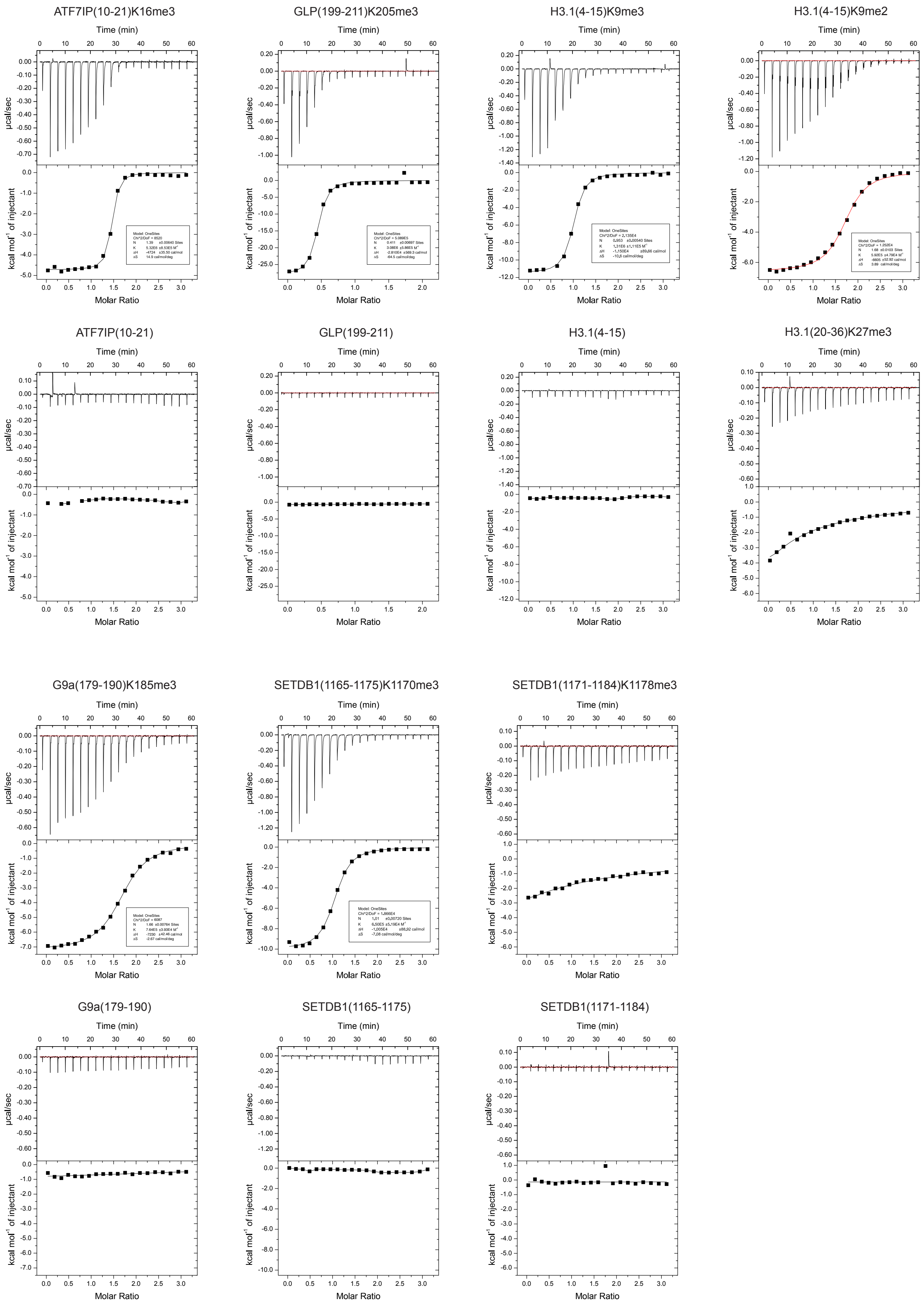

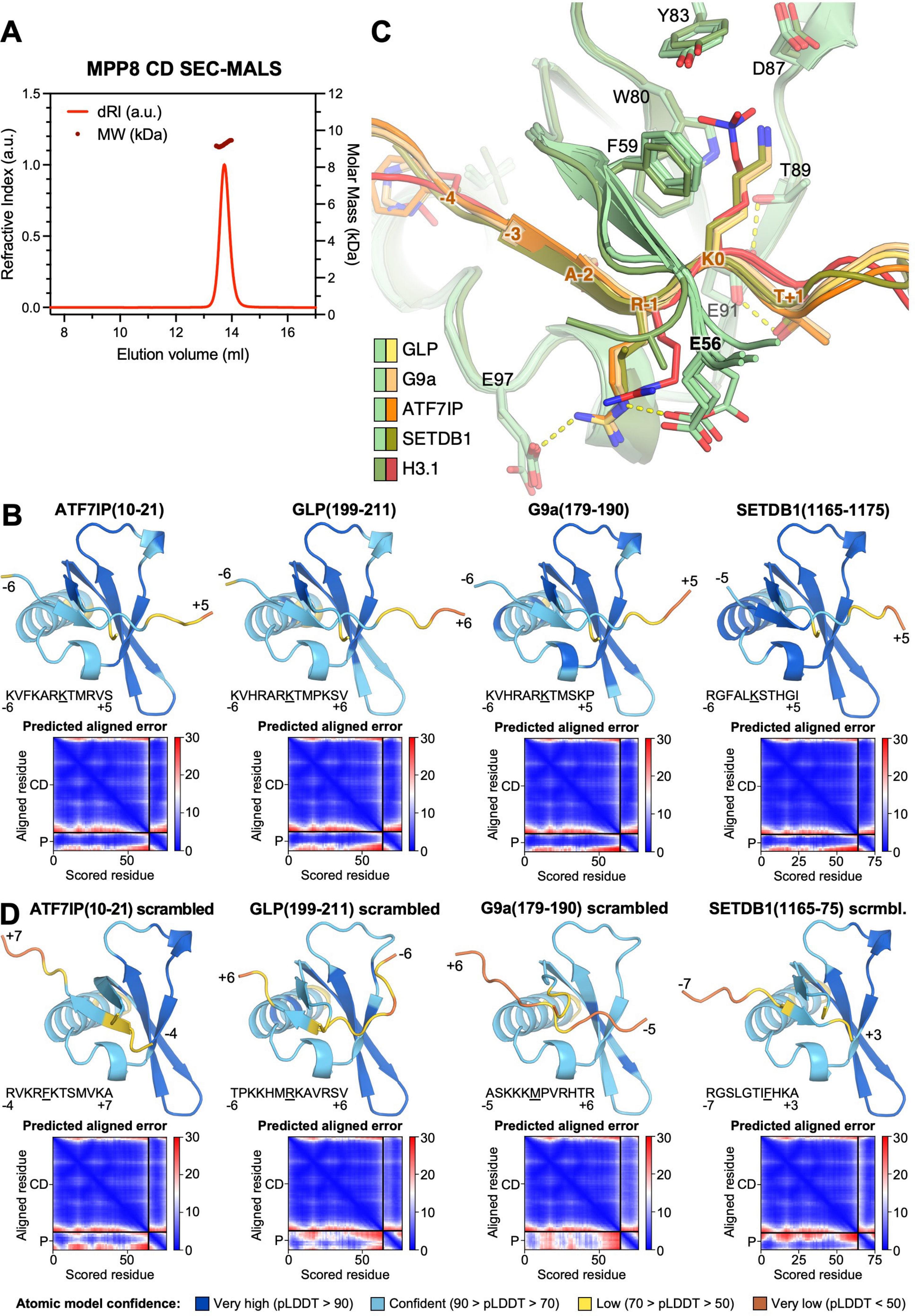
